# Single-Cell RNA Sequencing Defines Developmental Progression and Reproductive Transitions of *Pneumocystis carinii*

**DOI:** 10.1101/2025.04.21.649879

**Authors:** Aaron W. Albee, Steven G. Sayson, Alan Ashbaugh, Aleksey Porollo, George Smulian, Melanie T. Cushion

## Abstract

*Pneumocystis* species are host-obligate fungal pathogens that cause severe pneumonia in immunocompromised individuals. Despite their clinical importance, their life cycle remains poorly understood, in part because *Pneumocystis* depends on the host environment for most nutrients and requires sexual reproduction for survival, which occurs exclusively *in vivo*. This study presents the first single-cell RNA sequencing (scRNA-seq) atlas of *Pneumocystis carinii*, generated from isolated organisms recovered from the bronchoalveolar lavage fluid of infected rats to map the life cycle of *P. carinii*. Transcriptomes from 87,716 cells were analyzed using the 10x Genomics platform, revealing 13 transcriptionally distinct clusters representing key developmental stages, including biosynthetically active trophic forms, mating-competent intermediates, and asci undergoing sporulation. These states were characterized by expression of MAPK signaling components, β-glucan-modifying enzymes, and spore-associated genes, respectively. The scRNA-seq data supports previous evidence that these host-obligate fungi undergo sexual reproduction and provide new insights into the gene expression patterns associated with different life cycle phases. Biomarkers associated with ascus formation identified by scRNA-seq were validated by RT-qPCR, showing decreased expression levels in ascus-depleted populations treated with anidulafungin, a drug that halts ascus formation. More broadly, this approach provides a strategy for studying the full life cycles of fungal pathogens that cannot be continuously cultured.

**Importance:** *Pneumocystis* species *(spp.)* are clinically significant fungal pathogens that cannot be sustainably cultured *in vitro* due to their host obligate nature. This longstanding limitation has impeded progress in understanding their life cycle and identifying therapeutic vulnerabilities. Here, we apply scRNA-seq to *P. carinii* isolated directly from infected rat lungs, generating the first transcriptional map of its developmental progression. Our results define discrete gene expression states associated with trophic growth, mating activation, and ascus formation, and provide transcriptional evidence for a structured life cycle clarifying key developmental transitions and identifying potential regulatory targets for therapeutic intervention. Importantly, this study demonstrates that scRNA-seq can resolve the developmental biology of host-restricted fungal pathogens that cannot be cultured *in vitro*. This approach offers a generalizable framework for investigating other unculturable or obligate microbial pathogens directly within their native host environments, where traditional experimental tools are limited.

## Introduction

*Pneumocystis* species (spp.) are host-obligate fungal pathogens that cause life-threatening pneumonia in immunocompromised individuals. The human-specific species, *Pneumocystis jirovecii*, is responsible for *P. jirovecii* pneumonia (PjP) in patients with HIV/AIDS, organ transplants, or hematologic malignancies (1–3). These fungi adhere to type I pneumocytes in the alveolar lining, where their proliferation and the associated inflammatory response can lead to acute respiratory distress syndrome, ARDS (4–6). Despite their clinical importance, the basic biology of *Pneumocystis* spp., including reproduction, transmission, and persistence in the lungs, remains poorly understood.

*Pneumocystis spp.* lack an environmental stage and cannot survive without the host lung (7) and this constraint has made long-term culture and genetic manipulation unfeasible (8). Their mostly strict host specificity further complicates research; for example, *P. jirovecii* infects only humans, precluding direct study. To overcome this limitation, related species such as *P. carinii* (in rats) and *P. murina* (in mice) serve as experimental models for investigating infection dynamics, host-pathogen interactions, and life cycle progression (9). Genomic analyses show that *Pneumocystis spp.* have lost numerous biosynthetic and metabolic pathways common to most fungi (10, 11), including those for amino acid, carbohydrate metabolism, and lipid synthesis. These fungi complete their entire life cycle within the mammalian lung, relying on the host for nutrients and undergoing sexual reproduction *in situ.* As such, their biology dictates both their pathogenic strategy and the tools available for their study.

The life cycle has been largely determined by photomicrographs, which indicated an asexual phase via binary fission and a sexual phase resulting in the production of asci, structures essential for transmission to a new host (12, 13). Prior work from our group has shown that *Pneumocystis spp*. depend on sexual reproduction for survival and transmission (14, 15). Ascus formation, a key step in the sexual cycle, is required for completion of the life cycle and is inhibited by β-1,3-glucan synthesis inhibitors, such as anidulafungin. A better understanding of stage-specific gene expression is essential for identifying developmental regulators and potential vulnerabilities in the organism’s life cycle.

This study defines the first single-cell transcriptomic map of *P. carinii*, resolving gene expression patterns across its life cycle. It addresses three major challenges in *Pneumocystis spp.* research: the inability to culture the organism *ex vivo*, the essential role of sexual reproduction in its biology, and the lack of molecular detail regarding the sequence and coordination of life cycle stages. By adapting and optimizing a protocol for fungal cell isolation from bronchoalveolar lavage fluid (BALF), we preserved the transcriptomic profiles of individual cells, offering a first glimpse of *P. carinii* development within the host. This stage-resolved transcriptional framework lays the groundwork for targeted investigations of developmental regulation, therapeutic disruption of transmission, and comparative studies of other host-adapted fungal pathogens.

## Results

### Optimization of *Pneumocystis carinii* single-cell preparation for scRNA-seq

To facilitate single-cell transcriptomic profiling, a preparation protocol for *P. carinii* was developed to be compatible with the 10x Genomics platform (Supplemental Fig. S1A). Given that *P. carinii* populations are composed primarily of trophic forms, ∼90%, and only ∼10% asci (16), density gradient centrifugation was employed to enrich for asci (Supplemental Fig. S1B) (17) and to reduce host cell contamination, a common problem in *P. carinii* isolations. Organisms were obtained by bronchoalveolar lavage rather than lung homogenization, which further reduced the number of host cells. Post-enrichment viability consistently exceeded 80% (Supplemental Fig. S1C). Lysis conditions were adjusted by adding zymolyase to improve disruption of the 1,3-β-D-glucan-rich ascus wall (Supplemental Fig. S1D) (18). Two preliminary experiments informed adjustments to ensure sufficient sequencing depth for downstream analysis (Supplementary Tables ST1-ST3).

### Single-cell RNA-seq reveals 13 distinct transcriptional states across the *Pneumocystis carinii* life cycle

scRNA-seq of *P. carinii* isolated from bronchoalveolar lavage fluid (BALF) of infected rat lungs identified 13 transcriptionally distinct clusters (C1-C13), each representing a specific stage in the fungal life cycle (Fig. 1A). Pseudotime trajectory analysis placed clusters C1-C8 at earlier stages and clusters C9–C13 at later stages (Fig. 1B, Fig. 1C), with pseudotime values transitioning from low in C1-C8 to high in C9-C13. The heatmap of gene expression (Fig. 1D) shows the top 5 highly expressed genes in each cluster, with regulation indicated by a color gradient: upregulated genes in pink, downregulated genes in blue, and non-differentially expressed genes in white. GO term enrichment analysis (Fig. 2A) further supported these findings, revealing distinct biological processes associated with each cluster and showing three potential life cycle phases, described in detail below. Together, these data show a structured progression through the *P. carinii* life cycle, with transitions between trophic, mating, and sexual reproduction states.

**Figure 1.**
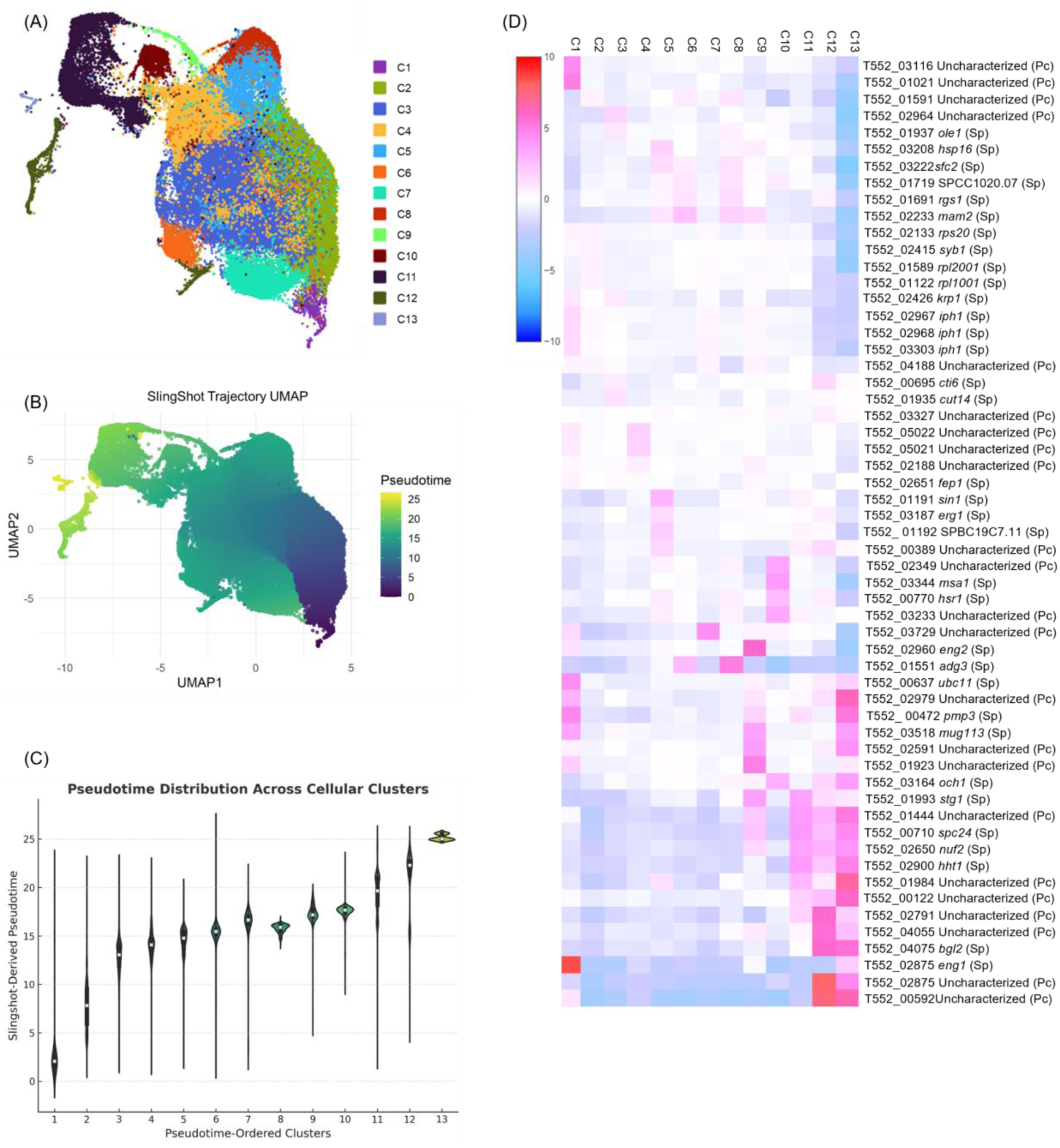
scRNA sequencing identifies transcriptionally distinct clusters in *P. carinii*. **Panel A:** Uniform Manifold Approximation and Projection (UMAP) of *P. carinii* cells, clustered into 13 transcriptionally distinct populations (C1-C13), indicated by color. **Panel B:** Pseudotime progression derived from Slingshot analysis, illustrating the inferred developmental trajectory from early clusters (dark purple) to late clusters (yellow).**Panel C:** Violin plot showing the distribution of Slingshot-derived pseudotime values across transcriptional clusters, ordered along the x-axis by developmental progression. Each violin represents cell density within a given cluster, with the median and interquartile range indicated by the embedded boxplots. **Panel D:** Heatmap showing the log-fold regulation of the top five differentially expressed genes (rows) across the 13 transcriptional clusters (columns). Each cell represents the average log-fold change in gene expression within each cluster.

**Figure 2.**
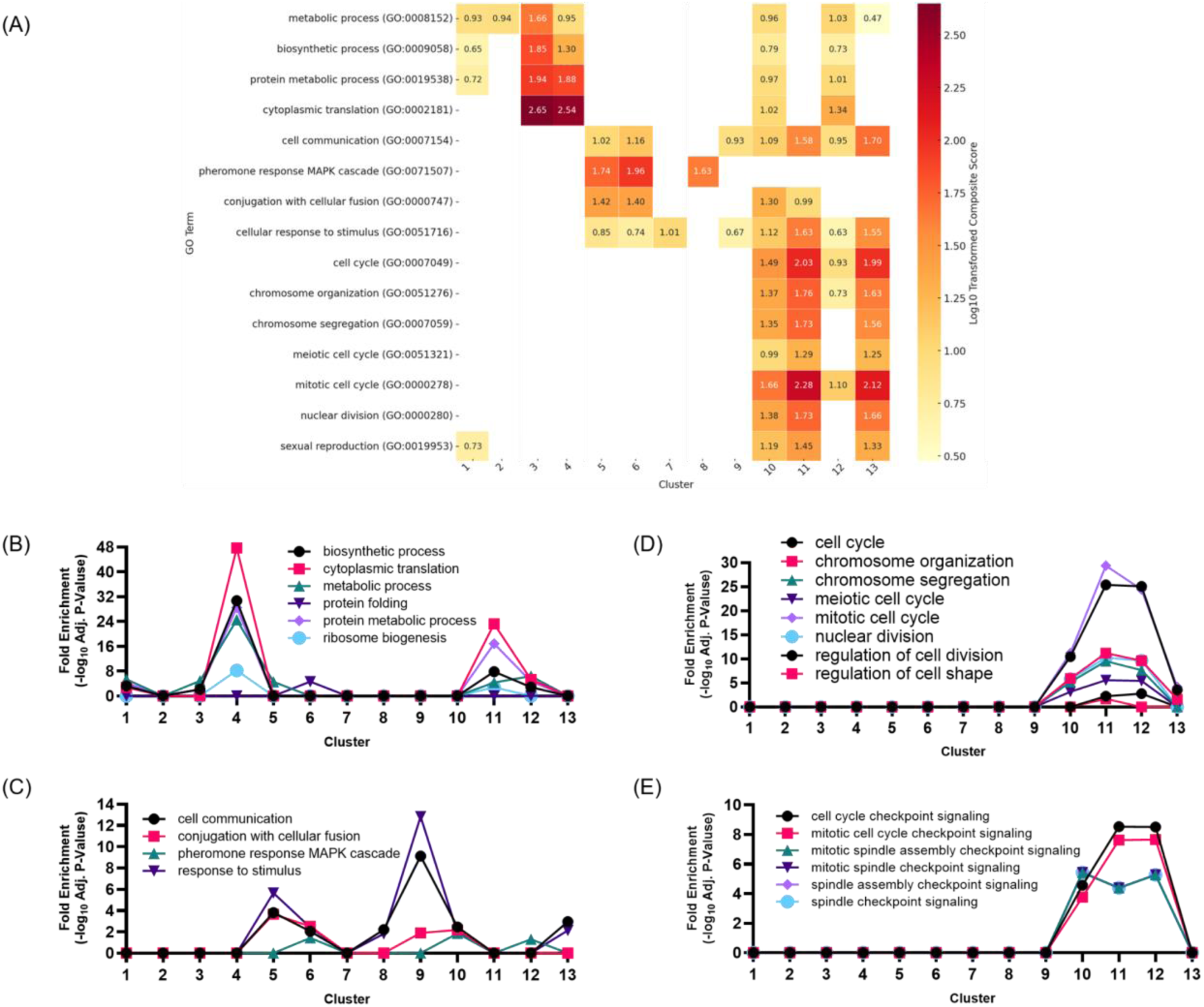
Transcriptional signatures from GO term enrichment define three distinct developmental transitions in *P. carinii*’s life cycle. **Panel A:** GO biological process enrichment heatmap showing 14 GO terms significantly enriched across transcriptional clusters. Enrichment is represented as a log_10_-transformed composite score (log_10_ fold change and adjusted p-value). Clusters are along the x-axis, and GO terms are listed on the y-axis. Dark red indicates higher enrichment (log_10_ fold change ≥ 2.5), while pale yellow indicates lower enrichment (∼0.5). **Panel B:** Line graph illustrating the enrichment of metabolic, biosynthetic, and protein metabolic processes across transcriptional clusters ordered by pseudotime. **Panel C:** Line graph showing the enrichment of sexual reproduction and signaling processes, including pheromone response and the MAPK cascade, across the transcriptional clusters. **Panel D:** Line graph highlighting enrichment of cell cycle-related processes, such as chromosome organization, segregation, and nuclear division, in late-stage clusters. **Panel E:** Line graph showing the enrichment of checkpoint-related pathways (e.g., mitotic spindle checkpoint signaling), which regulate cell division, particularly in late-stage development.

Clusters C1–C4 correspond to early trophic stages characterized by elevated metabolic and translational activity, with little indication of mitotic division. In <u>Cluster 1</u>, upregulation of *ptf1* (phosphoric monoester hydrolase) and *ubc11* (ubiquitin conjugation) (Table 1) reflects active metabolic processes, including phosphate metabolism, signal transduction, metabolic regulation, and protein turnover. The genes *pmp3* (plasma membrane proteolipid involved in ion homeostasis) and *mug113* (meiotic regulator) likely represent remnants of expression from Cluster C13, linking early and late stages of the life cycle. Downregulation of the mitotic gene *stg1* (cell cycle regulation, G2/M) and ascus-related cell wall genes (*gas1*, *och1*) (Table 2) supports the suppression of mitosis and ascus-related genes upregulated in Cluster C11-C13. GO term enrichment (Table 3) highlights processes such as cytoplasmic translation, biosynthesis, and metabolic processes, suggesting a metabolically active state, while sexual reproduction-related pathways remain minimally expressed, likely reflecting residual mRNA from Cluster C13.

**Table 1:**
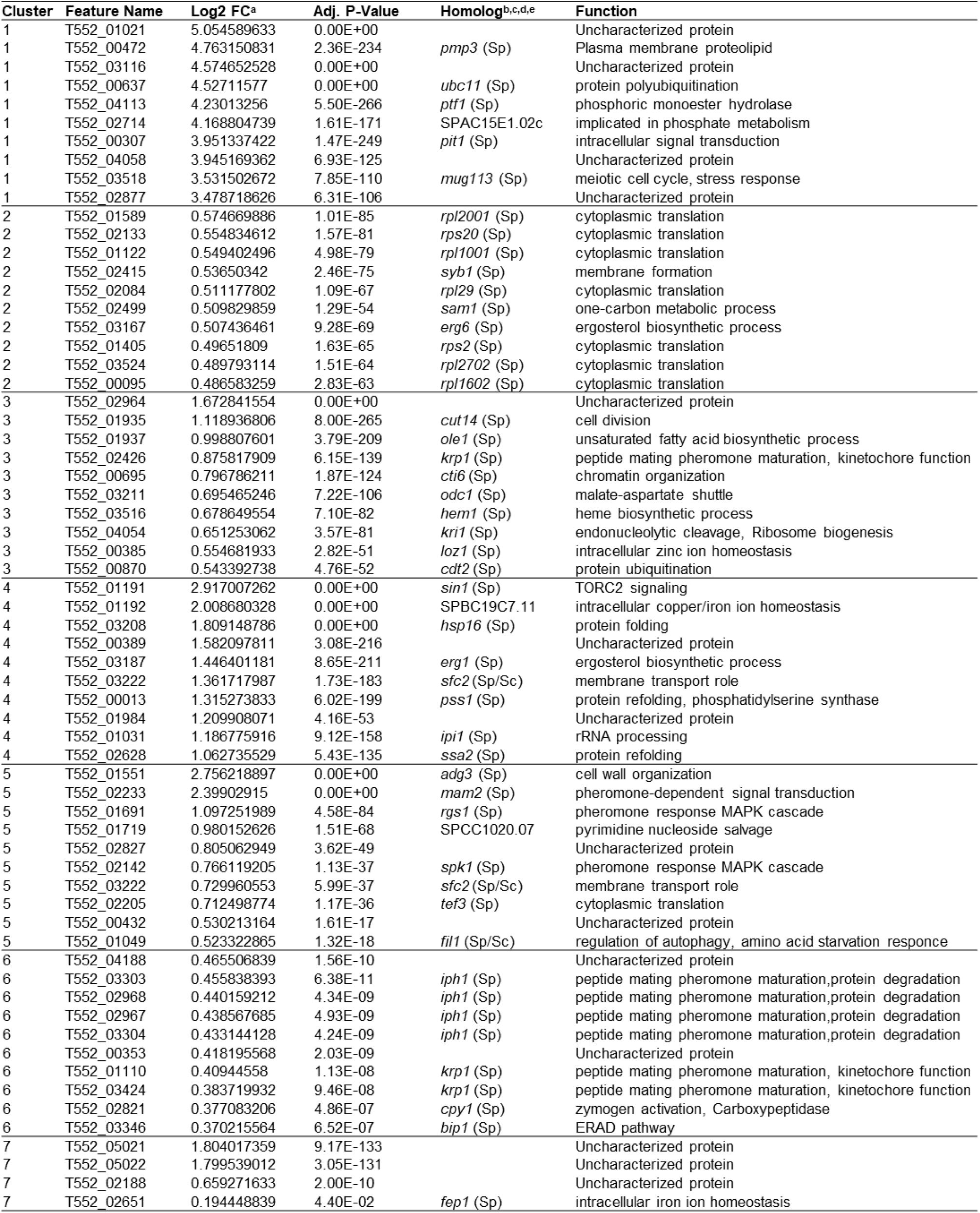
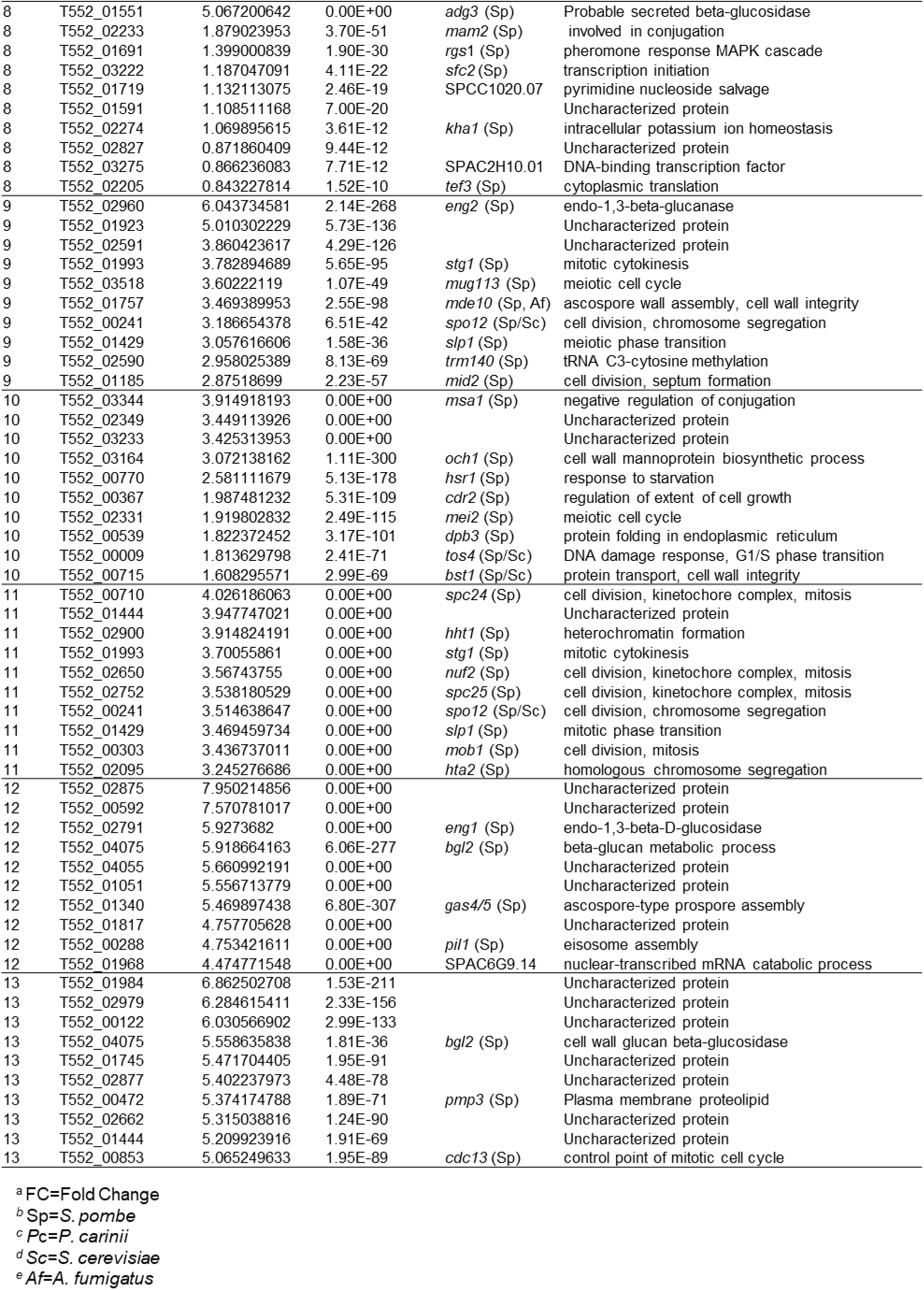
Top 10 Up-Regulated Genes in Each Cluster of *P. carinii*.

**Table 2:**
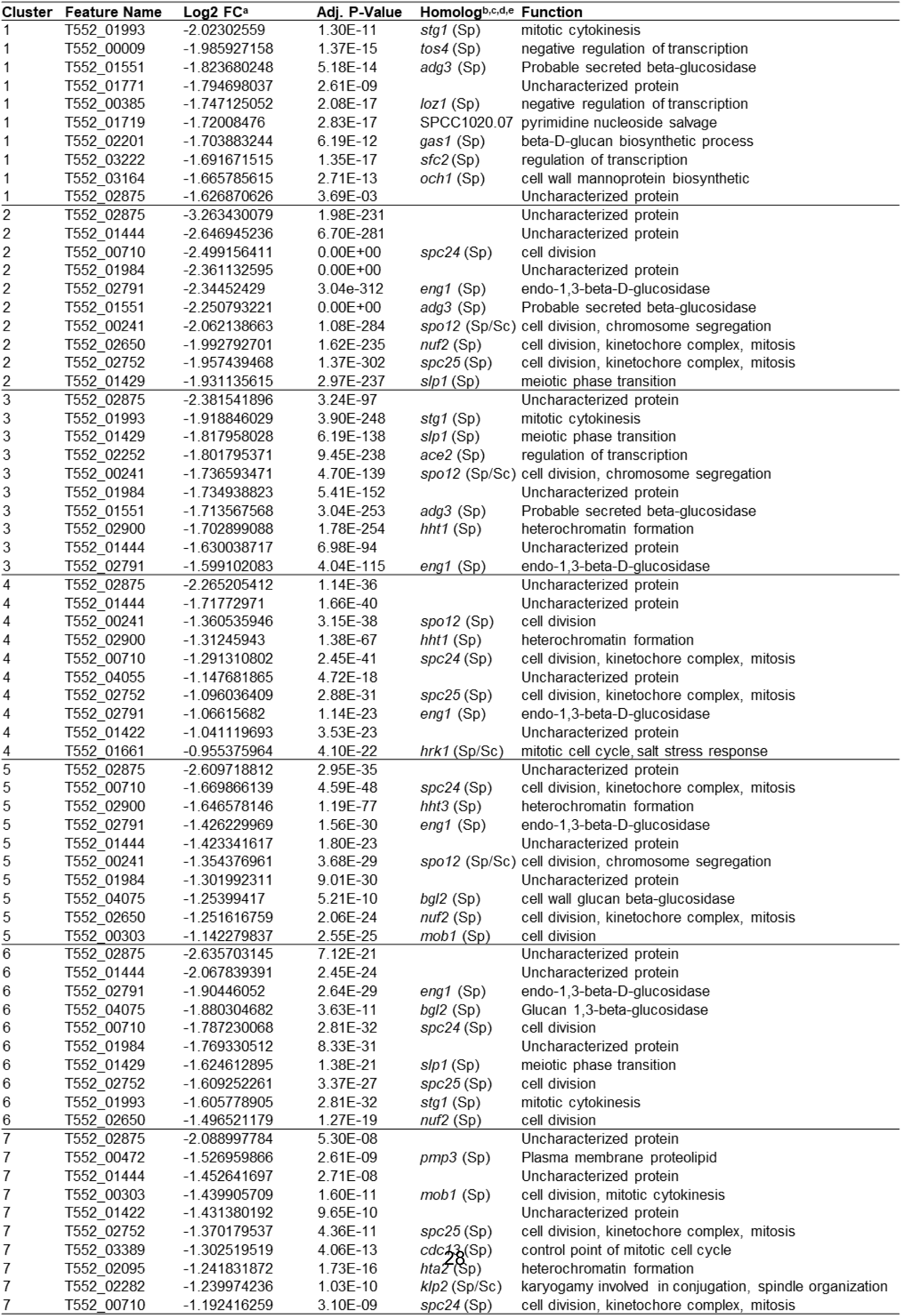
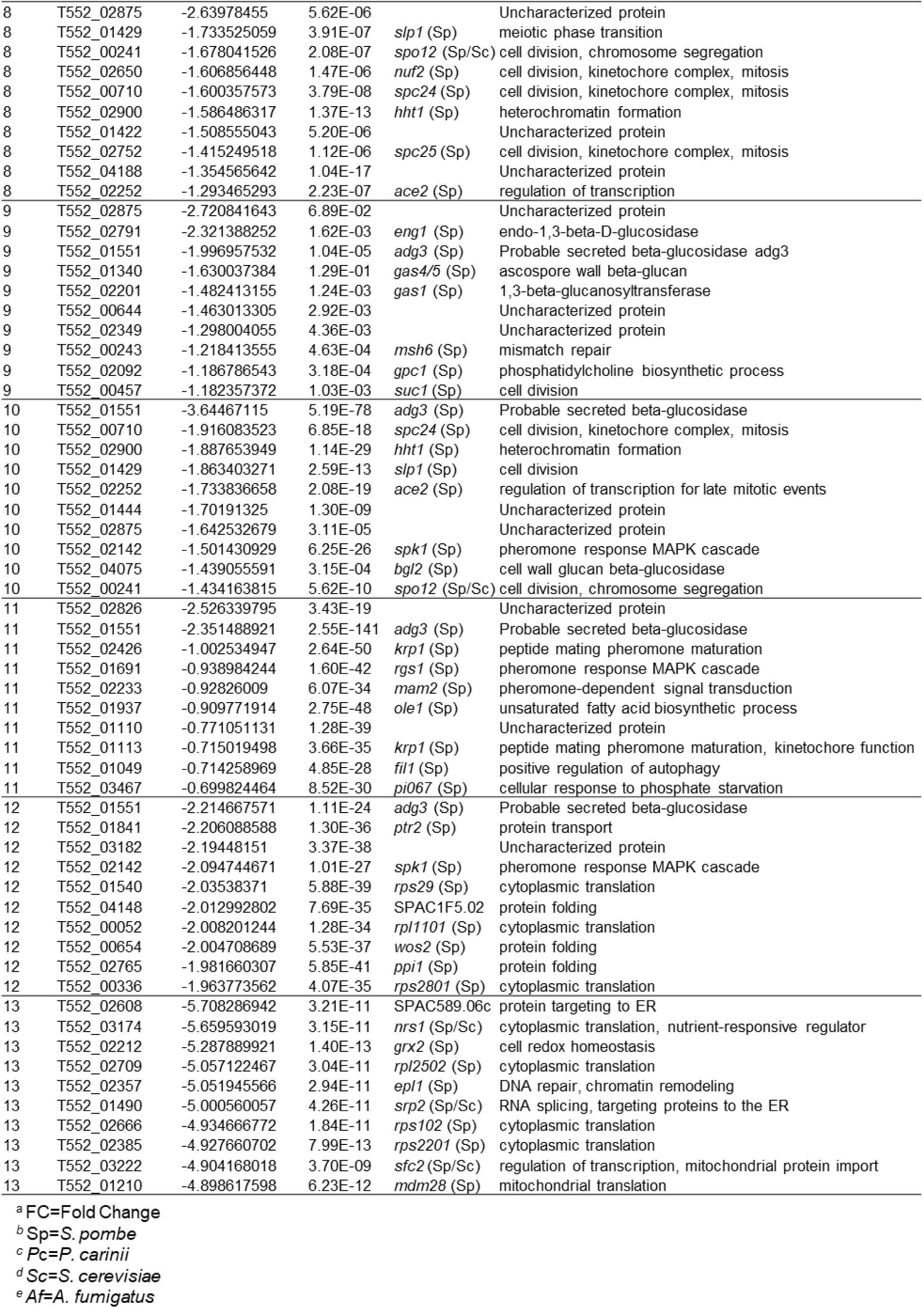
Top 10 Down-RegulatedGenes in Each Cluster of *P. carinii*.

**Table 3:** Enriched GO Terms Identified by G:Profiler Across scRNA-seq Clusters of *P. carinii:*

| Cluster | GO Term Name | GO Term ID | Enrichment Score | Adjusted P-Value |
| --- | --- | --- | --- | --- |
| 1 | structural constituent of ribosome | GO:0003735 | 39.56 | 2.73E-40 |
| 1 | metabolic process | GO:0008152 | 4.83 | 2.96E-25 |
| 1 | protein metabolic process | GO:0019538 | 2.18 | 3.26E-29 |
| 1 | biosynthetic process | GO:0009058 | 2.13 | 2.11E-31 |
| 1 | sexual reproduction | GO:0019953 | 1.71 | 1.51E-08 |
| 2 | binding | GO:0005488 | 9.12 | 1.77E-19 |
| 2 | molecular function | GO:0003674 | 8.45 | 2.05E-30 |
| 2 | biological process | GO:0008150 | 7.98 | 1.02E-29 |
| 2 | protein binding | GO:0005515 | 5.52 | 5.05E-23 |
| 2 | metabolic process | GO:0008152 | 4.54 | 2.96E-25 |
| 3 | cytoplasmic translation | GO:0002181 | 47.7 | 2.01E-48 |
| 3 | peptide biosynthetic process | GO:0043043 | 41.33 | 4.66E-42 |
| 3 | biosynthetic process | GO:0009058 | 30.68 | 2.11E-31 |
| 3 | metabolic process | GO:0008152 | 24.53 | 2.96E-25 |
| 3 | ribosome biogenesis | GO:0042254 | 8.22 | 6.07E-09 |
| 4 | cytoplasmic translation | GO:0002181 | 23.2 | 2.01E-48 |
| 4 | protein metabolic process | GO:0019538 | 16.88 | 3.26E-29 |
| 4 | biosynthetic process | GO:0009058 | 7.81 | 2.11E-31 |
| 4 | metabolic process | GO:0008152 | 4.36 | 2.96E-25 |
| 4 | ribosome biogenesis | GO:0042254 | 2.53 | 6.07E-09 |
| 5 | chaperone-mediated protein folding | GO:0061077 | 5.59 | 2.59E-06 |
| 5 | protein folding | GO:0006457 | 4.6 | 2.49E-05 |
| 5 | conjugation with cellular fusion | GO:0000747 | 2.53 | 2.22E-04 |
| 5 | cell communication | GO:0007154 | 2.06 | 1.73E-12 |
| 5 | cellular response to stimulus | GO:0051716 | 1.33 | 1.92E-13 |
| 6 | regulation of conjugation with cellular fusion | GO:0031137 | 3.41 | 3.89E-04 |
| 6 | pheromone response MAPK cascade | GO:0071507 | 3.33 | 1.43E-02 |
| 6 | intracellular signal transduction | GO:0035556 | 2.65 | 6.75E-13 |
| 6 | cell communication | GO:0007154 | 2.45 | 1.73E-12 |
| 6 | conjugation with cellular fusion | GO:0000747 | 2.19 | 2.22E-04 |
| 7 | molecular function | GO:0003674 | 4.8 | 2.05E-30 |
| 7 | RNA polymerase II sequence-specific DNA binding | GO:0000978 | 4.65 | 2.22E-05 |
| 7 | DNA-binding transcription factor | GO:0000981 | 4.58 | 2.61E-05 |
| 7 | cis-regulatory region sequence-specific DNA binding | GO:0000987 | 4.51 | 3.06E-05 |
| 7 | negative regulation of transcription | GO:0000122 | 1.3 | 6.93E-03 |
| 8 | DNA-binding transcription repressor activity | GO:0001227 | 2.81 | 1.54E-03 |
| 8 | cell communication | GO:0007154 | 2.45 | 1.73E-12 |
| 8 | conjugation with cellular fusion | GO:0000747 | 2.19 | 2.22E-04 |
| 8 | pheromone response MAPK cascade | GO:0071507 | 1.85 | 1.43E-02 |
| 8 | cellular response to stimulus | GO:0051716 | 1.34 | 1.92E-13 |
| 9 | molecular function | GO:0003674 | 9.64 | 2.05E-30 |
| 9 | biological process | GO:0008150 | 9.26 | 1.02E-29 |
| 9 | negative regulation of cellular process | GO:0048523 | 5.44 | 1.71E-10 |
| 9 | negative regulation of biological process | GO:0048519 | 5.22 | 3.92E-10 |
| 9 | cell communication | GO:0007154 | 2.2 | 1.73E-12 |
| 9 | cellular response to stimulus | GO:0051716 | 0.98 | 1.92E-13 |
| 10 | meiotic cell cycle | GO:0051321 | 11.21 | 2.68E-06 |
| 10 | metabolic process | GO:0008152 | 5.35 | 2.96E-25 |
| 10 | cellular response to stimulus | GO:0051716 | 5.06 | 1.92E-13 |
| 10 | sexual reproduction | GO:0019953 | 5.04 | 1.51E-08 |
| 10 | cell communication | GO:0007154 | 3.81 | 1.73E-12 |
| 11 | mitotic cell cycle | GO:0000278 | 29.39 | 4.09E-30 |
| 11 | cell cycle | GO:0007049 | 25.38 | 4.19E-26 |
| 11 | cellular response to stimulus | GO:0051716 | 12.72 | 1.92E-13 |
| 11 | nuclear division | GO:0000280 | 10.23 | 5.85E-11 |
| 11 | chromosome segregation | GO:0007059 | 9.54 | 2.85E-10 |
| 12 | cell cycle | GO:0007049 | 25.07 | 4.19E-26 |
| 12 | nuclear division | GO:0000280 | 9.69 | 5.85E-11 |
| 12 | chromosome segregation | GO:0007059 | 7.68 | 2.85E-10 |
| 12 | metabolic process | GO:0008152 | 6.42 | 2.96E-25 |
| 12 | mitotic cell cycle | GO:0051321 | 5.45 | 2.68E-06 |
| 13 | mitotic cell cycle | GO:0000278 | 24.42 | 4.09E-30 |
| 13 | cell communication | GO:0007154 | 11.76 | 1.73E-12 |
| 13 | cellular response to stimulus | GO:0051716 | 11.47 | 1.92E-13 |
| 13 | sexual reproduction | GO:0019953 | 6.69 | 1.51E-08 |
| 13 | meiotic cell cycle | GO:0051321 | 5.45 | 2.68E-06 |

In <u>Cluster 2</u>, several ribosomal proteins (Table 1) are upregulated, along with genes involved in protein synthesis, transportation, and cellular metabolism (*syb1*, *sam1*, *erg6*), reflecting a cellular state primed for biosynthetic activity and biomass accumulation. Concurrently, the downregulation of genes essential for mitotic progression and chromosome segregation (*spc24*, *spc25*, *nuf2*, *slp1*, *spo12*), as well as those involved in cell remodeling (*eng1*, *adg3*), suggests a transient suppression of cell division processes. Together, these expression patterns are consistent with translational readiness and metabolic investment over progression through mitosis or meiosis. GO term enrichment (Table 3) for translation and metabolic processes underlines the continued biosynthetic activity in these cells.

<u>Cluster 3</u> shows upregulation of genes involved in lipid metabolism (*ole1*), mitochondrial transport, and heme biosynthesis (*odc1*, *hem1*), ribosome biogenesis (*kri1*), and transcriptional and chromatin regulation (*cti6*, *loz1*, *cdt2*), representing a shift toward metabolic remodeling and preparatory gene expression (Table 1). Upregulation of *krp1* (pheromone precursor maturation) suggests a readiness for mating.

Although *cut14*, involved in mitosis, is upregulated, it also plays a role in chromatin organization, a function that can occur outside mitosis in response to stimuli. Simultaneously, the suppression of genes critical for mitotic exit and cytokinesis (*slp1*, *ace2*, *eng1*, *spo12*, *stg1*) (Table 2), as well as chromatin structure (*hht1*), suggests a transient delay or checkpoint in the cell cycle. The downregulation of mitotic genes is consistent with the absence of mitotic division, supporting the roles of *cut14* and *cti6* in chromatin structure outside mitosis. GO term enrichment (Table 3) highlights metabolic and biosynthetic processes, reinforcing the cluster’s focus on metabolism.

<u>Cluster 4</u> reflects a cellular adaptation to environmental or physiological stress, marked by upregulation of stress-response and membrane maintenance pathways and concurrent downregulation of mitotic progression and chromatin-related features (Table 2). Specifically, the increased expression of *sin1* and *hsp16*, along with chaperones like *ssa2*, suggests activation of pathways that protect protein integrity and membrane function. Simultaneously, the induction of lipid biosynthesis genes (*erg1*, *pss1*) and ribosome biogenesis factors (*ipi1*) indicates an investment in cellular repair and metabolic recalibration. In contrast, the downregulation of genes essential for chromatin structure (*hht1*), cell cycle progression (*spo12*, *spc24*, *spc25*), and cell wall remodeling (*eng1*) implies a deliberate slowing or arrest of cell division. This coordinated shift likely enables the cell to prioritize homeostasis and survival over proliferation, a strategic response to ensure long-term viability under suboptimal conditions. GO term enrichment (Table 3) for translation and biosynthesis reinforces the metabolically active state of these cells. In <u>Clusters 3 and 4</u>, the downregulation of *eng1* persisted while *adg3* was not downregulated in <u>Cluster 4</u>, suggesting the functions of these two genes may not be universally related.

GO terms were plotted in a line graph across the pseudotime-ordered clusters to illustrate the upregulation of biosynthetic and metabolic processes in the early clusters (Fig. 2B). The graph shows significant upregulation of these pathways in clusters C1-C4, with a marked peak in early stages, aligning with the high metabolic and translational activity observed in these clusters. As cells progress through pseudotime, there is a decline in the expression of these pathways, reflecting a shift away from biosynthesis and metabolism. A resurgence of metabolic and biosynthetic activity occurs in <u>Cluster 11</u> (Fig. 2B), just prior to the completion of sexual reproduction.

### Mating-competent trophic clusters (C5, C6, C8) show activation of pheromone signaling and repression of mitotic activity

Clusters 5, 6, and 8 reflect a transition toward a mating-competent state with activation of mating-related signaling and continued repression of mitotic activity. In <u>Cluster 5</u> (Table 1), upregulation of *mam2*, *rgs1*, and *spk1* indicates activation of pheromone signaling and mating pathways, likely in response to environmental cues or nutrient limitation. This is supported by increased expression of *fil1* and *adg3*, suggesting nutrient stress and metabolic remodeling. Elevated *tef3* and *sfc2* may aid in the synthesis and trafficking of mating-related proteins. The upregulation of *adg3* (putative β-glucosidase), although not directly linked to mating, occurs alongside mating genes. The enzyme *adg3* breaks down glucans outside the cell and may be involved in scavenging substrates during mating, preparing the cells for later β-glucan synthesis in Clusters 12 and 13. Downregulation of mitotic and chromosome segregation genes (*spc24*, *nuf2*, *spo12*, *mob1*), chromatin packaging (*hht3*), and cell wall remodeling (*eng1*, *bgl2*) indicates suppression of cell division, suggesting entry into the mating program. The continued downregulation of *eng1* across these clusters implies its specific role in ascus development. GO term enrichment for cell communication and pheromone response, MAPK signaling supports activation of mating-related processes (Table 3).

In <u>Cluster 6</u>, upregulation of multiple transcripts for pheromone maturation peptidases such as *iph1* and *krp1* (Table 1) is observed, while mitotic and cell cycle regulators remain repressed (Table 2). Downregulation of *bgl2* suggests that *adg3* activity is distinct from the regulated endo-1,3-β-D-glucosidase activity of *eng1* and *bgl2*. GO term enrichment for pheromone response and signal transduction reinforces this cluster’s focus on mating (Table 3).

<u>Cluster 8</u> shows upregulation of *mam2* and *rgs1* (Table 1), indicating pheromone response pathway activation. Concurrent expression of *adg3*, ion transporters (*kha1*), and trafficking-related genes (*sfc2*, *SPCC1020.07*, *SPAC2H10.01*) suggests resource reallocation for mating, possibly including membrane remodeling. Downregulation of core cell cycle and mitosis regulators (*slp1*, *spo12*, *nuf2*, *spc24*, *spc25*, *hht1*, *ace2*) indicates suppression of mitotic progression and chromosomal segregation, consistent with a G1 arrest or early meiotic entry. These transcriptional changes indicate a halt in proliferation to prioritize mating readiness. GO term enrichment for cell communication and pheromone response, MAPK signaling further supports mating activation (Table 3).

GO terms related to signaling, mating, and conjugation pathways were plotted across the pseudotime-ordered clusters to capture the dynamic regulation in <u>Clusters 5, 6, and 8</u> (Fig. 2C). A clear increase in these pathways is observed from <u>Cluster 5</u> to <u>Cluster 6</u>, followed by another peak in <u>Cluster 8,</u> indicating the activation of mating processes in these clusters. This pattern aligns with the upregulation of mating-related genes (*mam2* and *rgs1*)(Table 1), reinforcing the activation of mating signaling as the cells transition through these stages.

### Cluster 7 functions as a transcriptional checkpoint separating mating and meiosis before sexual reproduction

<u>Cluster 7</u> exhibits a unique transcriptional profile that marks a regulatory pause between mating and sexual reproduction. While genes involved in transcriptional regulation, such as *fep1* (iron uptake regulator), are upregulated (Table 1), no mating or meiotic markers are expressed in this cluster. The downregulation of genes related to cell division and mitotic regulation (*mob1*, *spc25*, *cdc13*, and *spc24*) (Table 2) suggests that <u>Cluster 7</u> functions as a pause point, halting the progression from mating to sexual reproduction. GO term enrichment (Table 3) reveals terms associated with transcriptional regulation and negative regulation of transcription, further supporting the role of <u>Cluster 7</u> in this regulatory pause. These findings are also reflected in the line graphs, which show the downregulation of metabolic, biosynthetic, mating signaling, and conjugation processes in Clusters 1–6, confirming the regulatory pause in Cluster 7 (Fig. 2B, Fig. 2C). This shift in gene expression highlights the paused state in <u>Cluster 7</u> before the cells proceed to sexual reproduction in subsequent clusters.

### Late Sexual Development (C9-C13) Shows Progression through Meiosis, Chromosome Segregation, and Ascus Wall Formation

<u>Clusters 9 to 13</u> represent the progression of sexual reproduction, with distinct gene expression patterns reflecting the transition from meiotic initiation to ascus formation. In <u>Cluster 9</u>, significant upregulation of meiotic genes (*stg1*, *mug113*, *spo12*, *slp1*, *mde10*, *mid2*) (Table 1) suggests entry into meiosis, including chromosome segregation, spore wall biosynthesis, and cytokinesis. Upregulation of *eng2* and *mde10*, involved in cell wall β-glucan remodeling, marks the onset of β-glucan metabolism; however, key genes for β-glucan biosynthesis remain absent, with crucial late-stage genes still downregulated.

Downregulation of mitotic cell wall enzymes (*eng1*, *gas1*, *gas4/5*), DNA repair factors (*msh6*), and cell cycle regulators (*suc1*, *gpc1*) indicates suppression of the vegetative cell cycle and mitotic checkpoints (Table 2), allowing a shift to meiosis. GO terms related to negative regulation of molecular function, cellular processes, and response to stimuli reflect coordination for sexual reproduction (Table 3).

<u>Cluster 10</u> continues meiosis with upregulation of *mei2*, *msa1*, and *tos4*, indicating meiotic initiation and regulation of gene expression. Upregulation of *cdr2*, *och1*, and *bst1* supports cell cycle remodeling, possibly to reorient polarity (Table 1). Downregulation of mitotic and chromatin regulators (*spc24*, *slp1*, *hht1*, *ace2*) (Table 2) confirms repression of mitosis and chromosome segregation, emphasizing meiotic commitment. GO term enrichment supports meiosis and sexual reproduction as cells progress toward ascus formation (Table 3).

<u>Cluster 11</u> shows upregulation of kinetochore genes (*spc24*, *nuf2*, *spc25*), histone genes (*hht1*, *hta2*), and mitotic exit regulators (*slp1*, *mob1*, *spo12*), indicating a shift toward robust mitotic proliferation after meiosis (Table 1). Downregulation of meiotic and mating-related genes (*krp1*, *rgs1*, *mam2*) (Table 2) supports the transition from meiosis to mitosis. GO term enrichment confirms this mitotic phase, marking the only mitosis observed in the life cycle. While downregulation of mating-related genes may be preventing mating inside the ascus.

<u>Cluster 12</u> reflects a physiological state focused on cell wall remodeling and environmental adaptation rather than proliferation. Upregulation of *eng1*, *bgl2*, *gas4/5*, and *pil1* indicates cell wall and membrane reorganization (Table 1). Concurrent downregulation of ribosomal genes (*rps29*, *rps2801*, *rpl1101*), protein-folding machinery (*wos2*, *ppi1*), and nutrient acquisition genes (*ptr2*) indicates suppressed growth and signaling pathways. This transcriptional program suggests a differentiated, non-proliferative state, consistent with ascus maturation. GO term enrichment for chromosome segregation and mitotic division supports this conclusion (Table 3).

<u>Cluster 13</u> shows upregulation of *bgl2* and *pmp3*, indicating cell wall and membrane preservation under stress, and possibly late-stage cell cycle arrest or quiescence in preparation for spore release. *Cdc13*, a telomere-capping protein, is upregulated, supporting genomic stability over replication. Downregulation of genes involved in redox homeostasis (*grx2*), nutrient-responsive cell cycle regulation (*nrs1*), and ribosomal proteins (*rps102*, *rps2201*, *rpl2502*) (Table 2) suggests suppression of translation and cell growth. Downregulation of *sfc2* and *mdm28* indicates reduced metabolic activity and organelle biogenesis. This gene expression pattern supports the hypothesis that *P. carinii* cells enter a quiescent, structurally stabilized state, likely a dormancy-like response to host stress or nutrient limitation. GO term enrichment highlights the final steps of sexual reproduction and ascus stress response (Table 3).

GO terms associated with sexual reproduction are illustrated in a line graph (Fig. 2D), demonstrating the dynamic upregulation of meiosis and mitosis across the ascus-related clusters <u>C10-C13</u>. A distinct peak in expression around <u>Clusters 12</u> and <u>13</u> marks the culmination of ascus formation. Similarly, GO terms associated with meiosis and mitosis checkpoint signaling exhibit sharp peaks between <u>Clusters 10</u> and <u>13</u>, confirming the regulation of meiotic and mitotic divisions across these stages (Fig. 2E). The previously mentioned resurgence of biosynthetic and metabolic pathways (Fig. 2B) tapers down to baseline in <u>Cluster 13</u>.

### RT-qPCR Confirms Ascus-Specific Expression of Late-Stage Marker Genes Identified by Single-Cell RNA Sequencing in *Pneumocystis carinii*

To validate stage-specific gene expression patterns identified by scRNA-seq, reverse transcription quantitative PCR (RT-qPCR) was performed on *P. carinii* RNA isolated from rats treated with anidulafungin, a β-1,3-D-glucan synthase inhibitor that halts ascus production. The inhibition of ascus formation was expected to reduce or eliminate the expression of late-stage ascus markers. RNA from treated animals was compared to RNA from untreated controls containing all life cycle stages. Expression of three scRNA-seq-defined late-stage marker genes, *T552_01968* (*bgl2*), *T552_01043* (*dmc1*), and *T552_01932* (*msa1*) was measured (Fig. 3A). The late-stage markers were chosen because, while genes in clusters C1–C8 have more shared functions, the marker genes for C12 and C13 are more specific to these clusters, making them ideal for validating stages.

**Figure 3.**
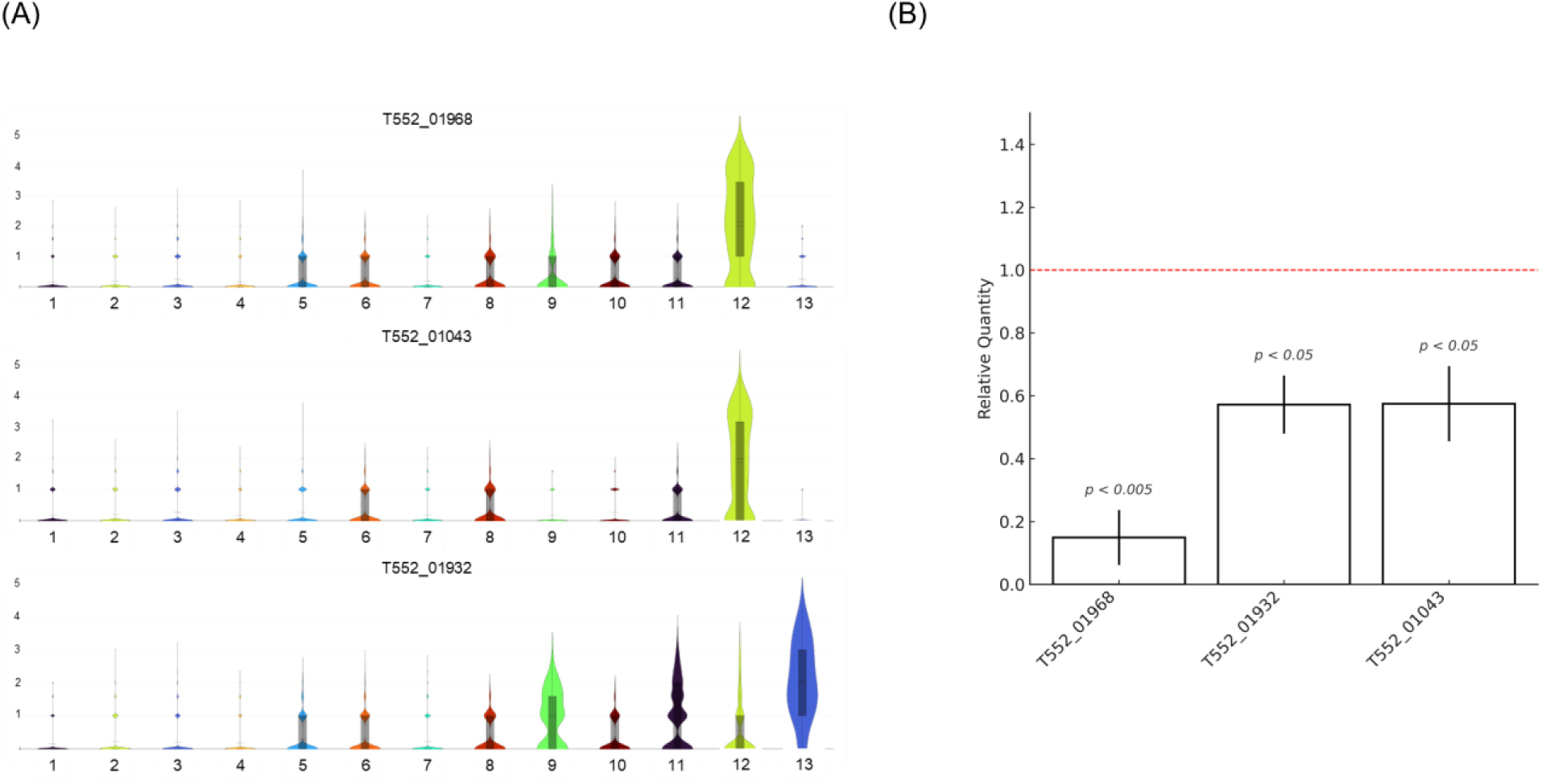
Validation of scRNA-seq-defined ascus marker gene expression by RT-qPCR. **Panel A:** Violin plots of the expression patterns of three ascus-associated marker genes (T552_01968, T552_01932, and T552_01043) across 13 transcriptional clusters (C1-C13) identified by scRNA sequencing. These genes are strongly enriched in Clusters C12 and C13, consistent with their roles as late-stage ascus markers. **Panel B:** Bar graph showing RT-qPCR validation of these genes in RNA isolated from ascus-depleted *P. carinii*-infected rats treated with anidulafungin for 3 weeks (zero asci verified by microscopic evaluation), compared to untreated controls (all life cycle stages represented, verified by microscopic evaluation). Expression of all genes is significantly reduced in the ascus-depleted group (p < 0.05 to p < 0.01), with relative quantities ranging from ∼0.05 to 0.80. The red dashed line indicates baseline expression in untreated animals. Error bars represent the standard error of the mean (SEM).

**Figure 4.**
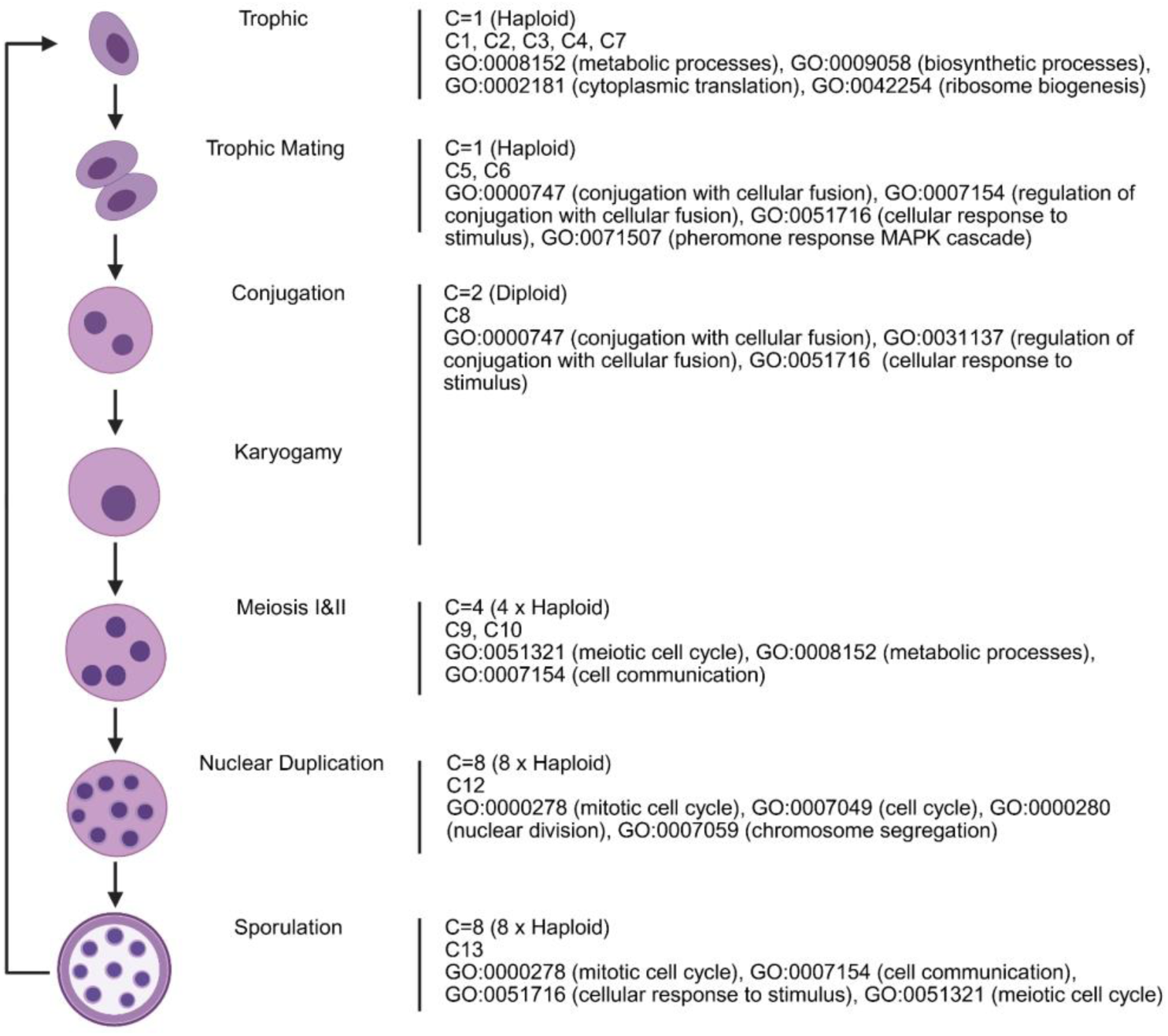
The life cycle of *P. carinii* from scRNA-seq trajectory analysis and GO enrichment. The figure illustrates the progression of the *P. carinii* life cycle, highlighting transcriptional clusters (C1–C13), associated ploidy levels (haploid, diploid, tetraploid, octoploid), and enriched gene ontology (GO) terms. Early clusters (C1-C5) represent biosynthetically active trophic forms (1C, haploid), characterized by upregulation of translation, ribosome biogenesis, and metabolic processes. Clusters C5 and C6 correspond to mating-competent haploid trophic cells, enriched for signaling and pheromone-related genes. Cluster C8, representing conjugation (2C, diploid), exhibits peak transcriptional complexity and marks sexual commitment. Karyogamy follows in C9, while C10 and C11 represent meiotic divisions (4C, tetraploid), enriched for meiotic and cell cycle processes. Cluster C12 corresponds to nuclear duplication (8C, octoploid), and C13 defines mature ascospore-containing asci, enriched for sporulation-related genes. This model suggests coordinated progression from trophic growth to sexual differentiation and ascus formation.

**Figure 5.**
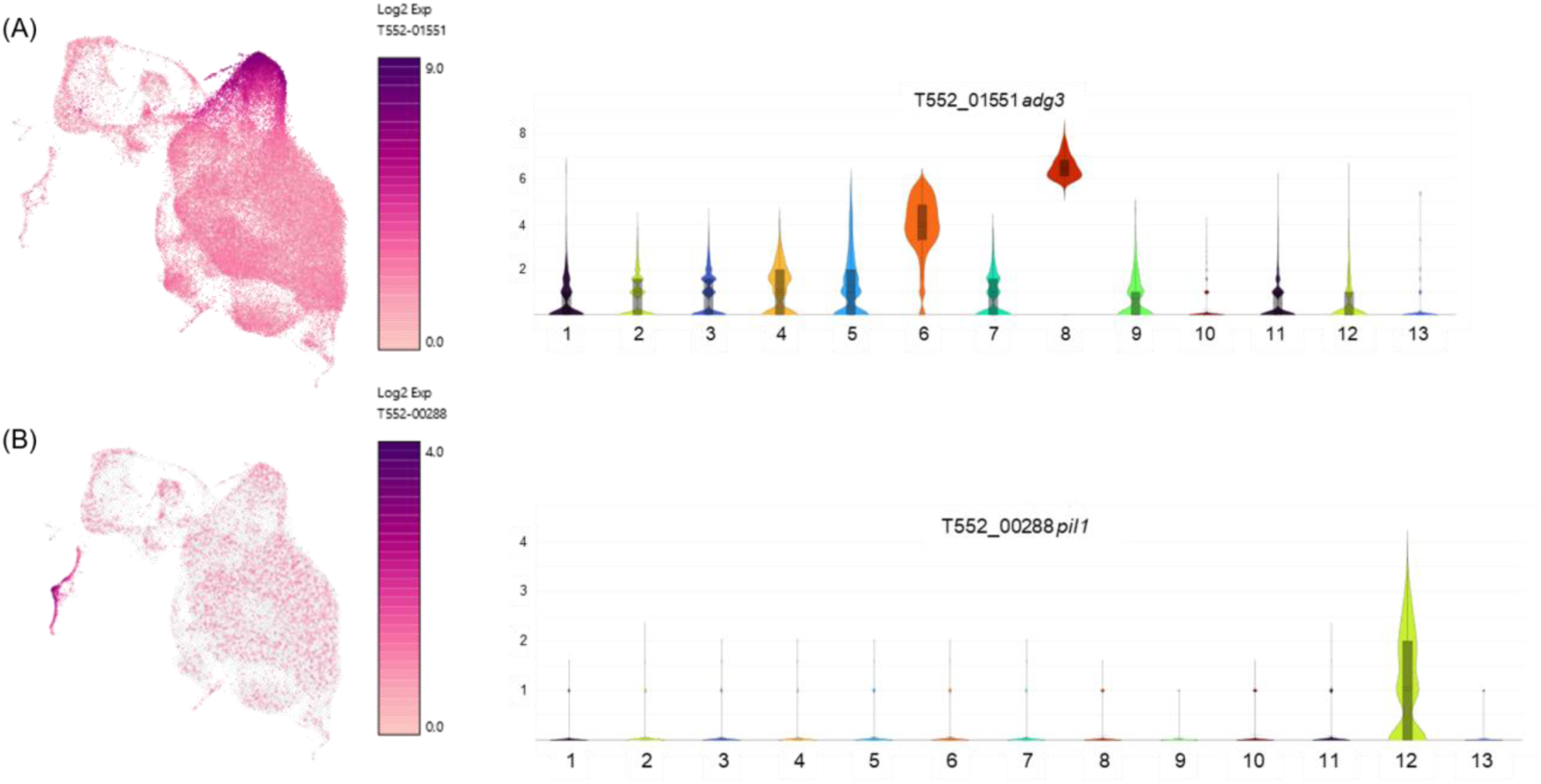
Expression dynamics of *adg3* and *pil1* during transcriptional transitions from trophic growth to ascus maturation in *P. carinii*. **Panel A:** UMAP feature plot (left) and violin plot (right) of *adg3* (T552_01551), a β-glucosidase gene predicted to encode a surface-localized hydrolase. *adg3* is moderately expressed in trophic clusters (C1-C4), sharply upregulated in mating-associated clusters (C5-C8), and repressed in reproductive stages (C10-C13). This expression pattern suggests roles in nutrient acquisition and cell wall remodeling during mating, with downregulation coinciding with ascus wall stabilization. **Panel B**: UMAP feature plot (left) and violin plot (right) of *pil1* (T552_00288), a homolog of eisosome-associated membrane organizers involved in ascospore formation. *pil1* expression is specific to clusters C12 and C13, corresponding to stages of nuclear duplication and sporulation. These findings align with the role of Pil1 in membrane reorganization during ascospore encapsulation in other ascomycetes. Together, these gene expression profiles illustrate coordinated transcriptional transitions from trophic proliferation (C1-C4), through mating and fusion (C5-C8), to reproductive differentiation and ascus maturation (C9-C13).

All three late-stage marker genes exhibited significantly reduced expression in the ascus-depleted group relative to untreated controls (Fig. 3B). Relative quantities ranged from 0.15 for *bgl2* to 0.57 for *msa1* and *dmc1*, with all differences reaching statistical significance (p = 0.0035–0.0254). These reductions align with the restricted expression of these genes to <u>Clusters C12</u> and <u>C13</u> in the single-cell dataset (Fig. 3A), supporting their association with asci and late-stage development.

These results independently corroborate the scRNA-seq-defined late-stage ascus markers, demonstrating that transcripts enriched in late transcriptional clusters are selectively diminished following pharmacologic blockade of ascus formation. The combined single-cell and qPCR data support a developmental model in which *P. carinii* progresses through discrete transcriptional states that culminate in ascus maturation. Together, these findings establish a transcriptional map for dissecting *in vivo* developmental progression in *P. carinii*.

## Discussion

This study provides the first single-cell transcriptional map of *Pneumocystis carinii*, revealing a coordinated life cycle progression from early trophic growth to mating, meiosis, and ascus formation. Using scRNA-seq, we identified 13 transcriptionally distinct clusters, which align with morphologically inferred stages, offering a comprehensive model for *P. carinii* life cycle progression in its host-dependent environment. Our findings challenge previous models that proposed asexual replication of trophic forms (19), as mitosis is repressed in all trophic clusters (C1-C8), suggesting obligate sexual reproduction. The early trophic clusters (C1-C4, C7) are enriched for metabolic genes, underscoring their focus on growth and nutrient acquisition rather than replication, consistent with observations in other fungi such as *Candida albicans* and *Cryptococcus neoformans* (20–22). In the mating-associated clusters (C5-C8), upregulation of *mam2*, the pheromone receptor, marks the transition to sexual differentiation (23), with dynamic regulation of *mam2* and *map3* differing from previous genomic studies, which anticipated concomitant expression of mating genes. Additionally, *adg3*, a β-glucosidase upregulated during mating, suggests its role in cell surface remodeling and nutrient acquisition, separate from other β-glucan genes (24–26).

Clusters C9-C13 represent sexual reproduction stages, with genes involved in meiosis, mitosis, and sporulation, including *pil1*, which marks late-stage ascus formation (27–29). Unlike *S. pombe*, where *eng2* is involved in spore release (30), *P. carinii* may utilize *eng1* for septal ring dissolution, a potential spore escape mechanism. These findings reflect the evolutionary divergence of *P. carinii* from related fungi, likely driven by its host obligate life cycle. Despite these advancements, the potential underrepresentation of adherent trophic forms (13) lost during bronchoalveolar lavage fluid (BALF) collection remains a limitation, and future studies should optimize sampling methods to capture these forms, as they likely exhibit distinct transcriptional profiles.

In comparison with previously hypothesized life cycles, our results support key aspects of traditional models, including the dominance of trophic forms and obligate sexual reproduction leading to ascus formation, consistent with early electron microscopy studies. However, we provide novel insights that challenge earlier models. The absence of asexual replication in trophic forms, supported by the lack of mitotic gene expression, contrasts with the notion of binary fission proposed by earlier studies (31). Our findings also align with genomic studies suggesting primary homothallism, as all cells possess a single mating type locus, differing from earlier assumptions of heterothallism (10, 32). Furthermore, the identification of a transcriptional checkpoint in Cluster 7, a regulatory pause between mating and meiosis, has not been previously described. The sequential and distinct transcriptional states we identify further challenge the notion of overlapping processes suggested in earlier models (33, 34), providing a more refined understanding of the life cycle progression. Finally, the upregulation of nutrient scavenging genes during mating, consistent with fungal sexual induction under starvation conditions, adds a novel layer to our understanding of metabolic reprogramming during sexual differentiation (21). This study, therefore, refines and expands traditional life cycle models of *Pneumocystis*, supporting an obligate sexual reproduction model driven by primary homothallism and a tightly regulated developmental progression.

Despite these advancements, limitations include the potential underrepresentation of adherent trophic forms lost during bronchoalveolar lavage fluid (BALF) collection (13). Future studies should optimize sampling methods to capture *P. carinii* adherent to host pneumocytes, as these forms likely exhibit distinct transcriptional profiles. Expanding this approach to other species, such as *P. murina*, or using BALF from *P. jirovecii*, will further elucidate host adaptation strategies across *Pneumocystis* species. Overall, this study establishes a foundation for studying *P. carinii* and other obligate fungal pathogens in their native host environments using single-cell transcriptomics.

## Materials and Methods

### Animal Model for *Pneumocystis carinii* Pneumonia

To induce *Pneumocystis carinii* pneumonia, male Sprague-Dawley rats (Charles River, Wilmington, MA), weighing 125 to 150 grams, were administered 20 mg/kg of Depo-Medrol (Pfizer, New York, NY) subcutaneously once per week for eight weeks to induce immunosuppression. After two weeks of continuous immunosuppression, rats were inoculated intranasally with 1 × 10⁶ *P. carinii* organisms suspended in water (9).

### Anidulafungin Treatment for Depletion of Asci

Immunosuppression was maintained for 5 weeks post-inoculation. Rats were divided into two groups (n = 6 per group). One group received the anidulafungin (Eraxis, 5 mg/kg/day)(Pfizer, NY, NY) by intraperitoneal injection for an additional 3 weeks. The second group received no antifungal treatment. At the end of the 3-week treatment period (8 weeks post-inoculation), all animals were euthanized, and the lungs were harvested. Microscopic enumeration of *P. carinii* cysts and trophic forms was performed to confirm ascus depletion in the anidulafungin-treated group, as detailed in prior work by our lab (15).

### Enumeration of *P. carinii* Trophic and Ascus Forms

*P. carinii* life cycle stages were quantified using established microscopy-based enumeration protocols. Three 10-µL aliquots were placed on microscope slides (Fisher Scientific, Pittsburgh, PA), air-dried, and heat-fixed. To distinguish developmental forms, asci were visualized using cresyl violet acetate (CVA; Sigma-Aldrich, St. Louis, MO), while total nuclei, including trophic and ascus forms, were stained using a modified Wright-Giemsa stain (Leuko-stat; Fisher Scientific, Pittsburgh, PA). Counts were averaged to determine the concentration and ratio of developmental stages (35). The final cell suspensions were normalized to a target concentration of 700 cells/µL for loading into the 10X Genomics single-cell RNA-seq assay.

### Isolation of *Pneumocystis carinii* from Rat Lungs

*P. carinii* was extracted from rat lungs via bronchoalveolar lavage (BAL). To remove host cells, BAL fluids from three individual rats were pooled and sequentially filtered through 70 µm and 40 µm pore filters (Fisher Scientific, Pittsburgh, PA) (36). The filtrates were centrifuged at 2500 x g to pellet fungal cells, which were resuspended in 0.85% ammonium chloride solution at 37°C for 10 minutes to lyse red blood cells. After centrifugation, the resulting pellets were maintained in RPMI 1640 medium (Fisher Scientific, Pittsburgh, PA) supplemented with 20% fetal bovine serum (Cytiva, Sweden AB), MEM Non-Essential Amino Acids (Gibco, Pittsburgh, PA), MEM Vitamin Solution (Gibco, Pittsburgh, PA), penicillin-streptomycin (10,000 µg/mL,10,000 µg/mL; Gibco, Pittsburgh, PA), and vancomycin (5 mg/mL; Fisher Scientific, Pittsburgh, PA) (8). Following extractions, incubation at 37°C for 30 minutes in T25 tissue culture flasks facilitated host cell adherence and enabled the enzymatic degradation of extracellular DNA using DNase I (Thermo Fisher Scientific, Waltham, MA). DNase I was applied at a concentration of 100 U/mL for effective DNA removal in cell culture systems (37). *P. carinii* organisms were collected from the supernatant, followed by centrifugation at 2500 x g to pellet and resuspended in 1 mL of 2% Ficoll-Hypaque before placement on a Ficoll gradient (38). We used a Ficoll-Hypaque gradient (Ficoll: Millipore, Billerica, MD; Hypaque: Sigma, St. Louis, MO). The gradient was prepared with Ficoll concentrations diluted in 16% sodium diatrizoate, ranging from 2% to 12% increments (17). Following centrifugation, each layer was collected, and the cells were rinsed with cold Dulbecco’s Phosphate-Buffered Saline (DPBS; Fisher Scientific, Pittsburgh, PA) and pelleted by centrifugation at 2500 × g. To minimize cell clumping, pellets were gently resuspended using a 20-gauge ball-tip gavage needle (Becton Dickinson & Co., Franklin Lakes, NJ) before final centrifugation at 2500 x g. The resulting samples were reconstituted in Mg²⁺ and Ca²-free RPMI 1640 medium (Fisher Scientific, Pittsburgh, PA) supplemented as described above.

### Single-Cell RNA Sequencing Data Processing and Analysis

Raw sequencing reads from scRNA-seq experiments were processed using Cell Ranger v9.0 (10x Genomics). Reads were aligned to the *Pneumocystis carinii* B80 reference genome with removal of Major surface glycoprotein genes (Accession: GCF_001477545.1). Genes detected in fewer than 10 cells were excluded, and cell-level quality control (QC) filtering was applied using the following thresholds: UMI per cell (<400), genes per cell (<200), and gene expression complexity (log10[genes per UMI] ≤ 0.8) to remove low-quality, doublet, or damaged cells that would result in technical noise. Expression data from all samples were merged and normalized using SCTransform and PCA using the top 50 principal components determined by an elbow plot (39). Clustering was conducted using the Louvain algorithm with a resolution parameter of 0.2, and visualization was performed using Uniform Manifold Approximation and Projection (UMAP) to represent transcriptional heterogeneity.

### Visualization and Differential Gene Expression Analysis Using Loupe Browser

scRNA-seq data from *P. carinii* were analyzed using Loupe Browser v8.1.1 (10x Genomics) visualization (40). LoupeR converted Seurat v5.2.0 objects into a compatible format (10x Genomics Software LoupeR, version 1.1.4). Data were projected using UMAP with clusters determined by Cell Ranger. Differential gene expression was assessed using the Significant Feature Comparison Analysis tool, selecting genes with an average occurrence of more than 1 count per barcoded spot, with expression values derived from log₂-transformed unique molecular identifiers (UMI) counts.

### Gene Annotation and Manual Curation of *Pneumocystis carinii* Genes

*P.* carinii genes were annotated by mapping homologs to *S. pombe* genes (41). A custom BLAST database was generated from *S. pombe* protein sequences (Accession: GCF_000002945.1), against which *P. carinii* proteins (Accession: GCF_001477545.1) were aligned (42). Protein sequence comparisons were conducted using BLASTp on the public Galaxy server (version 22.05, usegalaxy.org) (43, 44). The threshold of the expected value (E-value) was set to <0.0001 to ensure high-confidence alignments. The top three matches with the highest bit scores were considered for annotation.

### GO Term Enrichment Analysis and Visualization

Gene Ontology (GO) enrichment analysis was performed using g:Profiler (version e112_eg59_p19_25aa4782)(45, 46) and gene annotations were sourced from Ensembl (47) Fungi release. A significance threshold of adjusted *p* < 0.05 was applied, with multiple testing corrections performed using both the Bonferroni method and False Discovery Rate (FDR) adjustment.

GO term enrichment for biological and molecular processes was conducted using Cluster Profiler, with a background gene set derived from the *Pneumocystis carinii* B80 genome (Accession: GCF_001477545.1) (48). Statistical significance was determined using Fisher’s exact test with Benjamini-Hochberg correction (P<0.05). Enrichment scores were log_10_-transformed for visualization.

All visualization and statistical analysis were performed in Python 3.10 using a Google Colab environment (Notebook ID: GO_Pseudotime_ClusterViz_2024) (49, 50), Heatmaps were generated with Seaborn v0.11.2, and line plots representing –log_10_ adjusted *p*-values (p<0.05) were produced using Matplotlib v3.5.3 and SciPy v1.10.1 (51, 52).

### Trajectory Analysis

Trajectory analysis was conducted using Slingshot v2.14.0 to infer lineage structures within cell populations (53). Input data comprised UMAP components derived from Seurat-generated single-cell transcriptomic data. A minimum spanning tree (MST) was constructed to define global lineage relationships, and cell lineages were assigned using Slingshot’s clustering-based approach. The root node was manually selected based on the most transcriptionally distinct cluster. Lineage-specific gene expression trends were analyzed using generalized additive models (GAMs) to identify genes that are dynamically regulated along inferred trajectories. Cell transitions and lineage progression were visualized using UMAP embedding, with cells ordered by the trajectory analysis.

### Marker Gene Validation by RT-qPCR

Marker genes in *Pneumocystis carinii* were selected for validation based on differential expression identified through scRNA-seq analysis. Target genes were the top upregulated transcripts from Clusters 12 and 13. Thymidylate synthase (TS; T552_02292) (p=0.9286411) was used as the internal reference (59). Gene-specific primers (Supplementary Table S2) were tested with a no-template negative control. RNA was extracted from *P. carinii* organisms using the Direct-zol RNA Miniprep Kit (Zymo Research, Irvine, CA, USA). First-strand cDNA was synthesized from extracted RNA using SuperScript IV VILO Master Mix (Thermo Fisher Scientific, Waltham, MA, USA) and stored at –80°C.

RT-qPCR was performed using Applied Biosystems 7500 Real-Time PCR System (Applied Biosystems, Foster City, CA, USA). Each 20 μl reaction contained 1× PowerUp SYBR Green Master Mix (Thermo Fisher Scientific, Waltham, MA, USA) and 500 nM of each primer, run in biological and technical triplicate. The thermocycling protocol consisted of 50°C for 2 minutes, followed by 95°C for 2 minutes, and 40 cycles of 95°C for 15 seconds, 59°C for 15 seconds, and 72°C for 1 minute. Fluorescence was measured during the 72°C extension step and melt curve analysis was performed to verify amplification specificity. The threshold cycle (ΔCT) value between the validation gene and the TS gene was calculated by subtracting the average CT value of the reference gene from the average CT value of the validation gene. ΔΔCT values were calculated by subtracting the ΔCT value of the reference sample from the experimental sample. Relative quantity is shown as 2-ΔΔCT.

## Data Availability

scRNA-seq files have been deposited in NCBI under accession no. 25126083

## Acknowledgments

The National Institutes of Health (NIH) R01HL146266 (M.T.C.) and the U.S. Department of Veterans Affairs (VA) 1I01BX004441-01 (M.T.C.) supported this work. M.T.C. is a research career scientist at the Cincinnati Veterans Affairs Medical Center.

